# Impact of U2-type introns on splice site prediction in *Arabidopsis thaliana* using deep learning

**DOI:** 10.1101/2024.05.13.593811

**Authors:** Espoir Kabanga, Soeun Yun, Arnout Van Messem, Wesley De Neve

## Abstract

In this study, we investigate the impact of introns on the effectiveness of splice site prediction using deep learning models, focusing on *Arabidopsis thaliana*. We specifically utilize U2-type introns due to their ubiquity in plant genomes and the rich datasets available. We formulate two hypotheses: first, that short introns would lead to a higher effectiveness of splice site prediction than long introns due to reduced spatial complexity; and second, that sequences containing multiple introns would improve prediction effectiveness by providing a richer context for splicing events. Our findings indicate that (1) models trained on datasets with shorter introns consistently outperform those trained on datasets with longer introns, highlighting the importance of intron length in splice site prediction, and (2) models trained with datasets containing multiple introns per sequence demonstrate superior effectiveness over those trained with datasets containing a single intron per sequence. Furthermore, our findings not only align with the two hypotheses we put forward but also confirm existing observations from wet lab experiments regarding the impact of length of an intron and the number of introns present in a sequence on splice site prediction effectiveness, suggesting that our computational insights come with biological relevance.

**Author summary:** In this study, we explore how intron characteristics affect the effectiveness of splice site predictions in *Arabidopsis thaliana* using deep learning. In particular, focusing on U2-type introns due to their prevalence in plant genomes and their relevance for large-scale data analysis, we demonstrate that both the length of these introns and the number of introns present in a sequence substantially influence prediction outcomes. Our findings highlight that deep learning models trained on data with shorter introns or multiple introns per sequence produce better predictions, aligning with observations from wet lab experiments regarding the impact of intron length and the number of introns per sequences on splice site prediction effectiveness.

## Introduction

In eukaryotic organisms, during messenger RNA (mRNA) processing, introns are excised, and exons are joined together to form the mature mRNA, which serves as the template for protein synthesis [1]. This process, which is illustrated in Fig 1, is known as splicing. It is carried out by the spliceosome, a complex composed of RNA and protein molecules. The spliceosome recognizes specific sequences at the boundaries between introns and exons, known as splice sites, and removes the introns while ligating the exons together [2]. Splice sites include both donor and acceptor sites [3]. A donor splice site, positioned at the 5’ end of an intron, is characterized by the dinucleotide `GT`, signaling the beginning of the intron to be removed during RNA splicing. An acceptor splice site, found at the 3’ end of an intron, is marked by the dinucleotide `AG`, indicating the end of the intron to be excised. These sites play crucial roles in the accurate removal of introns and the subsequent joining of exons, facilitating the production of mature messenger RNA. Precise splice site prediction is a crucial element in gene expression analysis [4].

**Fig 1.**
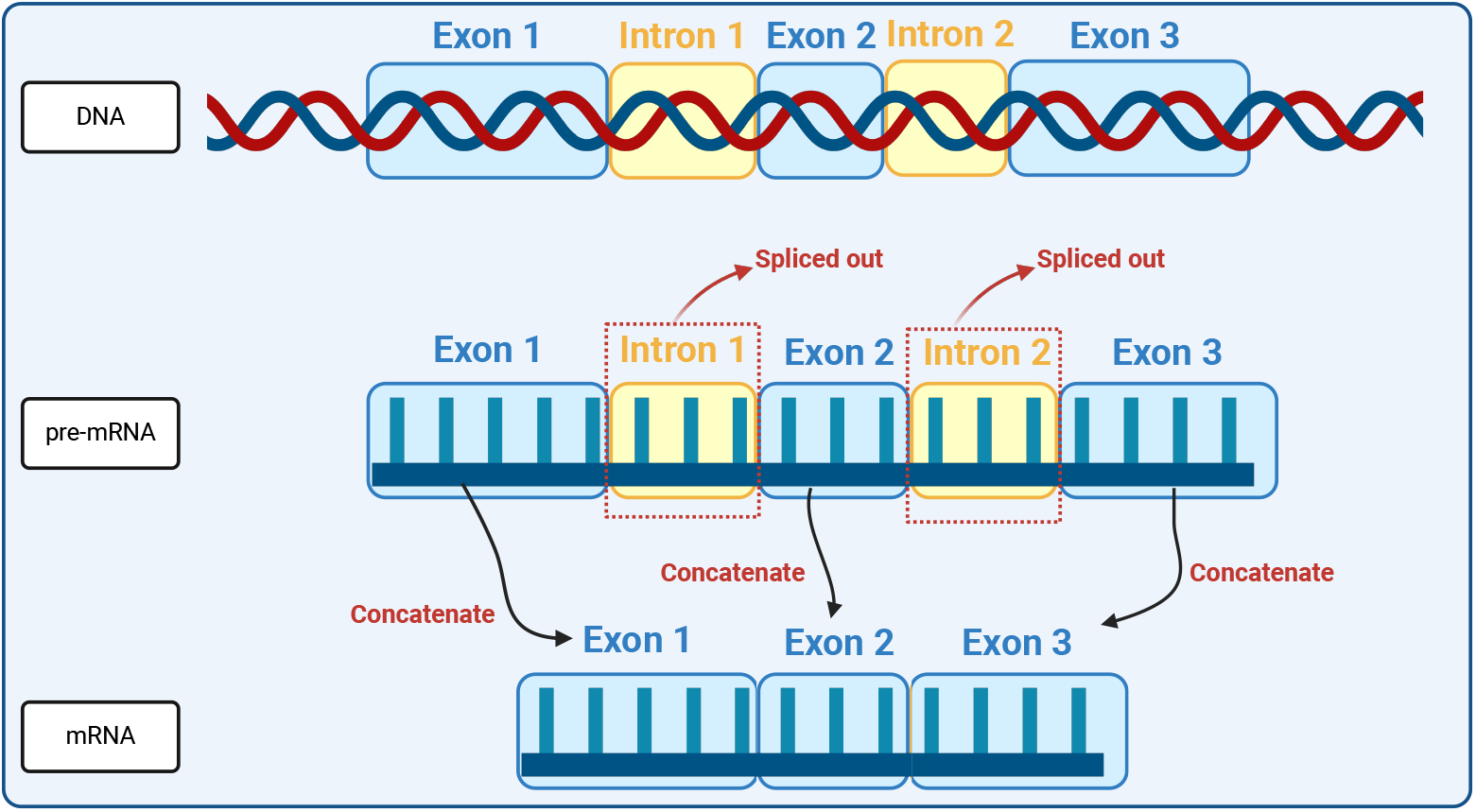
Splicing mechanism. Visualization of the process of RNA splicing where introns are removed from the pre-mRNA transcript and exons are joined together to form the mature mRNA sequence.

Introns can substantially affect gene expression in plants and many other eukaryotes in a variety of ways [5]. U12-type and U2-type introns are two distinct classes of introns found in eukaryotic genomes [6]. They differ in terms of their spliceosomal machinery and splicing mechanisms [7]. U12-type introns are a less common class of introns, constituting a small fraction of introns in most eukaryotic genomes. They are spliced out through a minor spliceosome, which is a smaller and less understood spliceosome complex compared to the major spliceosome. U12-type introns have distinct consensus sequences at the donor splice site and the branch point sequence. On the other hand, U2-type introns, defined in this study by `GT-AG` boundaries at their splice sites, are the most prevalent type of introns in eukaryotic genomes. They are spliced out through the major spliceosome, a larger and more common spliceosome complex. The major spliceosome recognizes a highly conserved intron-exon junction sequence at the donor splice site, as well as a branch point sequence near the acceptor splice site [8]. Our decision to focus on U2-type introns in the context of splice site prediction in *Arabidopsis thaliana* is driven by several factors. Primarily, the ubiquity of U2-type introns in plant genomes, including *Arabidopsis thaliana*, makes them an important subject for understanding the global splicing landscape and its regulatory mechanisms. Moreover, the vast amount of data available for U2-type introns facilitates the application of deep learning techniques, which require large datasets for effective model training. Additionally, the choice of *Arabidopsis thaliana*, a model organism in plant biology, over other species is particularly strategic due to its well-annotated genome and status as a benchmark in plant genetics and molecular biology research [9].

Deep learning has emerged as an important tool in computational biology, revolutionizing various aspects of bioinformatics [10], including the prediction of splice sites. The complex nature of gene splicing, coupled with the vast amount of genomic data available, necessitates advanced computational techniques for accurate prediction of splice sites. Deep learning models, for instance making use of convolutional neural networks (CNNs) [11–15], have shown remarkable proficiency in extracting complex features from DNA and RNA sequences, enabling them to discern subtle patterns crucial for splice site identification. However, the presence of U2-type introns, the predominant class of introns in eukaryotic genomes, poses unique challenges to splice site prediction [16] due to factors such as variability in their sequences [17], the presence of multiple potential splice sites or the complex regulatory elements associated with them [18]. Incorporating these distinctive characteristics into deep learning models increases their precision and generalizability when dealing with U2-type intron-related splice site recognition. This fusion of deep learning with the complexity of U2-type introns has the potential to refine our understanding of gene regulation mechanisms and facilitate more accurate gene expression analysis.

## Materials and methods

### Dataset

In this study, we use DNA sequences with donor splice sites and acceptor splice sites from *Arabidopsis thaliana*. This dataset was compiled and organized for SpliceMachine [19], where the authors used linear support vector machines (SVMs) with manually defined positional and compositional features to predict splice sites.

The dataset, which we will refer to as the SpliceMachine dataset, includes 9,175 positive samples (DNA sequences with true splice sites) and 264,507 negative samples (DNA sequences without true splice sites) for donor splice sites, and 9,278 positive and 277,255 negative samples for acceptor splice sites. Notably, each sequence in the dataset had a fixed length of 402 nucleotides, with the splice sites (`GT` and `AG` for donor splice sites and acceptor splice sites, respectively) precisely positioned in the middle of the sequence. Table 1 describes the characteristics of the dataset.

**Table 1.**
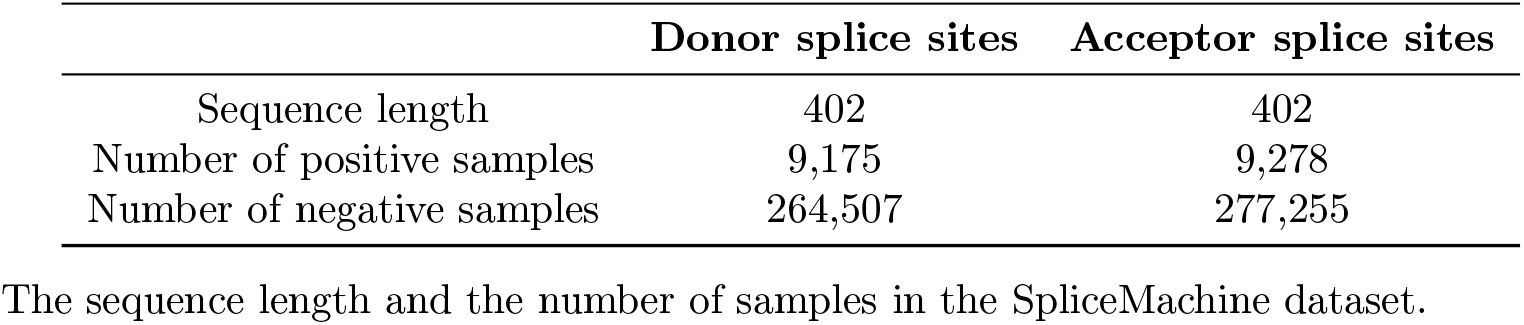
Characteristics of the SpliceMachine dataset.

To accurately identify U2-type introns, sequences were cross-referenced with the Intron Annotation and Orthology Database (IAOD) [20], which offers multi-species intron annotation. The combination of the SpliceMachine dataset and the IAOD dataset provided a solid framework for a comprehensive analysis of U2-type introns, ensuring accurate identification of splice sites. As described in Table 2, our analysis showed that the SpliceMachine dataset contained 7,811 sequences with U2-type introns for the donor splice site dataset, while the acceptor dataset had 7,970 such sequences. Of these, 7,571 unique U2-type introns were identified at 10,268 distinct locations in the donor dataset, indicating positional redundancy. Similarly, the acceptor dataset featured 7,621 unique U2-type introns at 10,454 unique locations.

**Table 2.** U2-type introns from IAOD found in the SpliceMachine dataset.

|  | Donor splice sites | Acceptor splice sites |
| --- | --- | --- |
| Number of U2-type introns present in the SpliceMachine dataset | 10,268 | 10,454 |
| DNA sequences from the SpliceMachine dataset with at least one U2-type intron | 7,811 | 7,970 |

The number of U2-type introns from IAOD found within the SpliceMachine dataset.

After filtering the dataset with sequences containing U2-type introns, we explored two key hypotheses related to splice site prediction models: (1) the impact of U2-type intron length and (2) the impact of the number of U2-type introns per sequence. Fig 2 provides an overview of these hypotheses and the overall data description.

**Fig 2.**
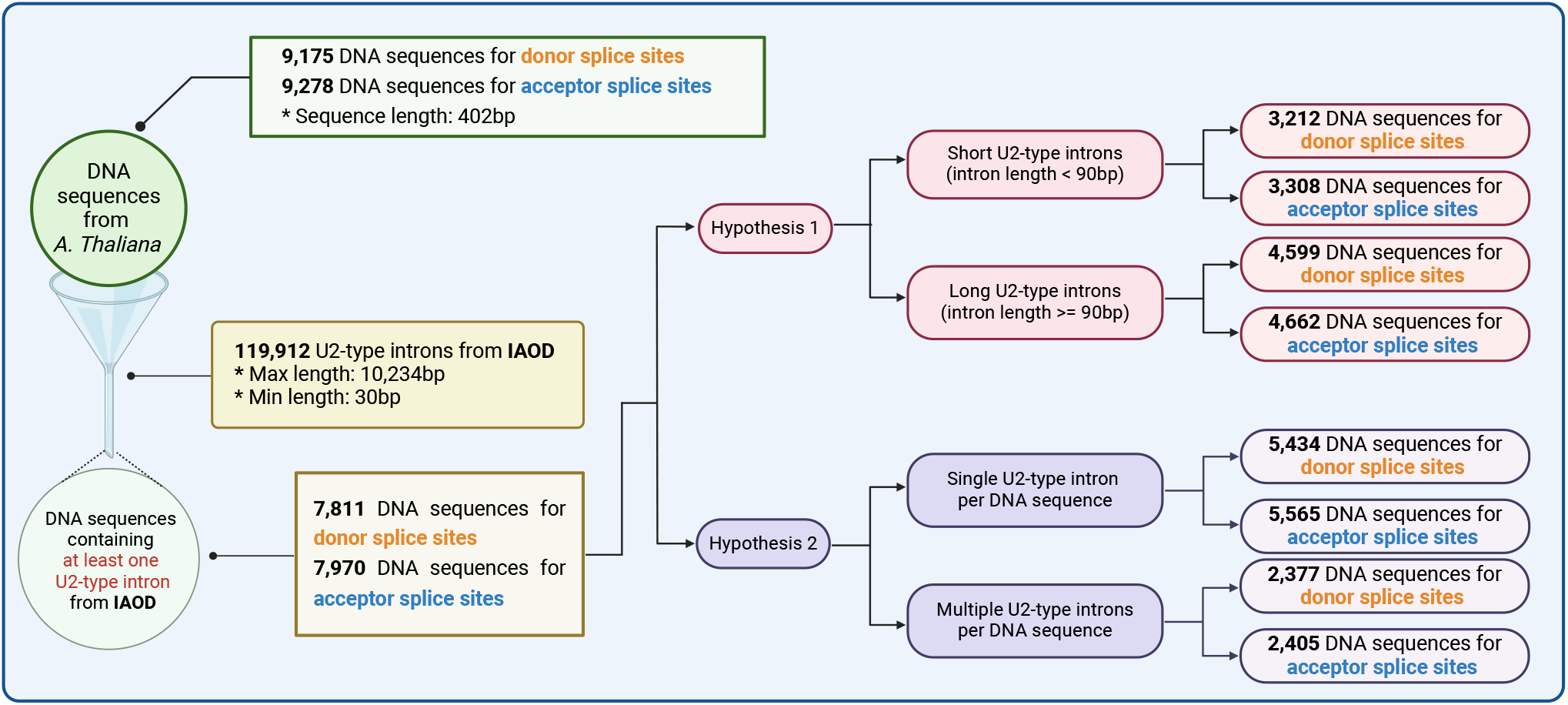
Hypotheses and dataset description. Overview of the two hypotheses explored in this study, along with a description of the dataset.

### Hypotheses

#### Hypothesis 1: Short introns lead to higher effectiveness in splice site prediction

Our first hypothesis revolves around the potential improvement in the effectiveness of splice site prediction models when trained with a dataset containing short introns, as opposed to long ones. This hypothesis is grounded in the complex molecular dynamics and biochemical factors inherent to RNA splicing.

The length of introns varies widely among different organisms and different genes. Generally, introns in animal genes are larger than those in plant genes. For example, human genes tend to have small exons separated by long introns with a mean and median length of 3356 and 1023 bp, respectively, whereas introns in *Arabidopis thaliana* have a mean and median length of 168 and 100 bp, respectively [21, 22]. Long introns can introduce spatial complexities that obstruct the precise alignment of intron-exon junctions during the splicing process. This spatial complexity, in turn, increases the likelihood of activating cryptic splice sites [23], which are sequences that can trigger unexpected splicing events.

Additionally, in cases with long flanking introns, the exon definition mechanism, which is the way the spliceosome identifies or recognizes exons to ensure they are properly included in the mature mRNA, becomes more dominant. This dominance presents a challenge to the spliceosome in accurately recognizing exon boundaries [24], as it relies on the precise identification of small exonic sequences, potentially resulting in exon-skipping events.

Given this molecular background, we anticipate that splice site prediction models will demonstrate superior effectiveness when applied to a dataset containing short U2-type introns. Short U2-type introns inherently possess fewer complexities, making them a promising focus for improving the effectiveness of splice site prediction.

Within the SpliceMachine dataset, we examined the U2-type intron length distribution. The donor set exhibited an average length of 102.69 bp, with a median length of 93 bp and a range from 59 to 202 bp. Similarly, the acceptor set had an average U2-type intron length of 102.43 bp, with the same median of 93 bp. However, the acceptor set had a broader range, with the length of U2-type introns varying from a minimum of 44 bp to a maximum of 202 bp. These observations highlight the importance of defining U2-type intron length within this context. Fig 3 shows the U2-type intron length distribution for both donor and acceptor datasets.

**Fig 3.**
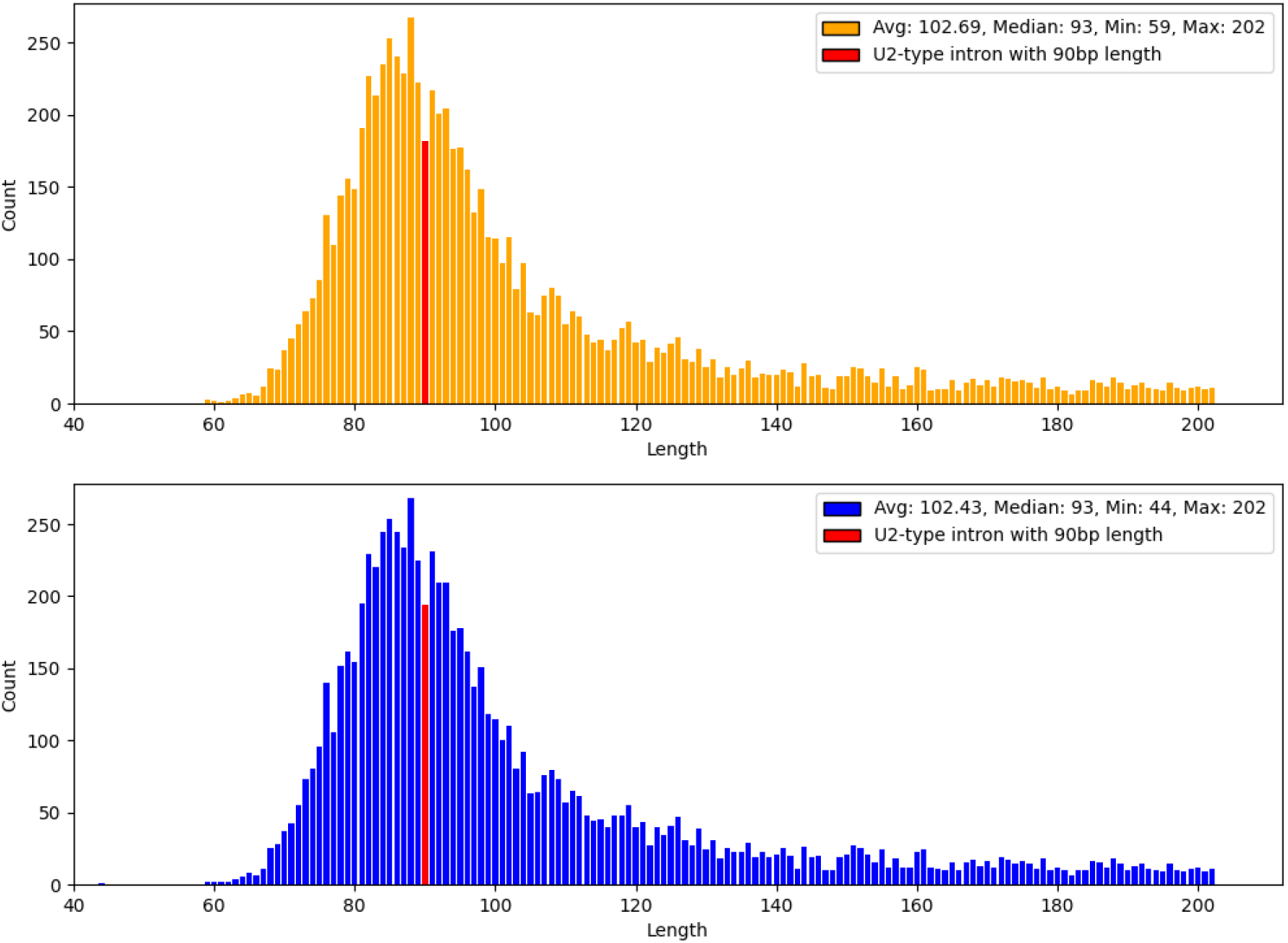
U2-type intron length distribution. The distribution of U2-type intron lengths for positive samples with donor splice sites (top) and for positive samples with acceptor splice sites (bottom). The red bar indicates a 90bp length; lengths to the left of this mark are categorized as short, while those to the right (including the 90 bp length) are categorized as long.

According to prior research, a mature intron, which consists of functional structure units that increase in number with the length of the intron and are linked to form a cohesive structure that interacts with mRNA, is generally not shorter than 80 bp [25]. This same study classifies introns into short introns (ranging from 0 to 120 bp) and long introns (exceeding 120 bp), with an almost even distribution between the two categories. Another study, specifically focusing on *Arabidopsis thaliana*, defines short introns as those shorter than 116 bp [26] and notes that the proportion of short introns falls within the range of 45% to 65%. In our study, we adopt the definition of U2-type introns shorter than 90 bp as short U2-type introns. This choice is supported by the consistent median U2-type intron length of 93 bp in both the donor and acceptor datasets.

Based on the aforementioned classification criterion, the SpliceMachine dataset included, on the one hand, 3,212 DNA sequences with short U2-type introns (41.1% of the positive samples) and 4,599 DNA sequences with long U2-type introns (58.9% of the positive sample) among sequences with donor splice sites. On the other hand, there were 3,308 DNA sequences with short U2-type introns (41.5% of the positive samples) and 4,662 DNA sequences with long U2-type introns (58.5% of the positive samples) among sequences with acceptor splice sites. These percentages confirm the validity of our 90 bp threshold for distinguishing between short and long U2-type introns [27].

To ensure a balanced and robust comparison, we organized the SpliceMachine dataset into six distinct subsets: a donor set with short U2-type introns, a donor set with long U2-type introns, a donor set with an equal mixture of short and long U2-type introns, an acceptor set with short U2-type introns, an acceptor set with long U2-type introns, and an acceptor set with an equal mixture of short and long U2-type introns. We randomly sub-sampled each of these six subsets in order to obtain six corresponding working sets of 3,212 samples. By maintaining an identical number of sequences in each subset, potential biases are minimized and any observed differences in prediction effectiveness can be attributed to the intrinsic properties of the sequences rather than to dataset size. To better handle the small number of positive samples, we performed sub-sampling of the negative samples to achieve a 1:5 ratio of positive to negative samples, resulting in 16,060 negative samples for each subset. This approach was to ensure a balanced representation of data to minimize bias. Finally, we randomly partitioned the data of each subset into 80% for training and 20% for testing.

#### Hypothesis 2: Multiple introns per sequence in a dataset lead to enhanced splice site prediction effectiveness

Our second hypothesis asserts that a dataset containing sequences with more than one U2-type intron will yield superior effectiveness in splice site prediction compared to a dataset containing sequences with only one U2-type intron. This hypothesis is built upon insights from splicing kinetics and the complex regulatory mechanisms controlling gene expression.

Specifically, kinetic analysis has indicated that the removal of the first intron in a sequence often improves the effectiveness with which subsequent splicing events occur. This improvement may arise from accelerated recruitment of the exon junction complex (EJC) to ligated exons or the establishment of a stable splicing framework that promotes the subsequent removal of introns [28].

Additionally, experimental data in [29] have shown that configurations of multiple introns within a transcript can efficiently influence the splicing machinery. For example, full-length long-read sequencing techniques have revealed that in most cases, upstream introns tend to be spliced before downstream ones. This order of splicing not only follows the transcriptional sequence but also indicates that early spliced introns may facilitate the processing of subsequent introns, potentially improving the effectiveness of splice site identification in multi-intron transcripts.

Consequently, this hypothesis anticipates that the effectiveness of splice site prediction models improves when they are trained on a dataset containing multiple U2-type introns per sequence.

We divided the sequences in the SpliceMachine dataset into two categories: one with sequences containing only one U2-type intron and one with sequences containing more than one U2-type intron. Specifically, in the case of donor splice sites, the dataset consisted of 5,434 DNA sequences with a single U2-type intron per sequence and 2,377 DNA sequences with multiple U2-type introns per sequence. Similarly, for acceptor splice sites, the dataset included 5,565 DNA sequences with a single U2-type intron per sequence and 2,405 DNA sequences with multiple U2-type introns per sequence.

We then organized the above two categories into six different subsets: a donor set with only one U2-type intron per sequence, a donor set with more than one U2-type intron per sequence, a donor set with an equal mixture of sequences with only one U2-type intron and sequences with more than one U2-type intron, an acceptor set with only one U2-type intron per sequence, an acceptor set with more than one U2-type intron per sequence, and an acceptor set with an equal mixture of sequences with only one U2-type intron and sequences with more than one U2-type intron.

As for the first hypothesis, we randomly sub-sampled each of these six subsets in order to obtain six corresponding working sets of 2,377 samples. To better handle the small number of positive samples, we performed sub-sampling of the negative samples to achieve a 1:5 ratio of positive to negative samples, resulting in 11,885 negative samples for each subset. Following this, we randomly partitioned the data of each subset into 80% for training and 20% for testing.

### Model description

In this study, whose workflow is described in Fig 4, we built a novel CNN model named IntSplicer (Intron Splicer). To evaluate the effectiveness of our model, we benchmarked it against three existing models: SpliceRover, SpliceFinder, and DeepSplicer. Each model has its unique architecture and approach to splice site prediction, thus making it possible to perform a comprehensive comparison.

**Fig 4.**
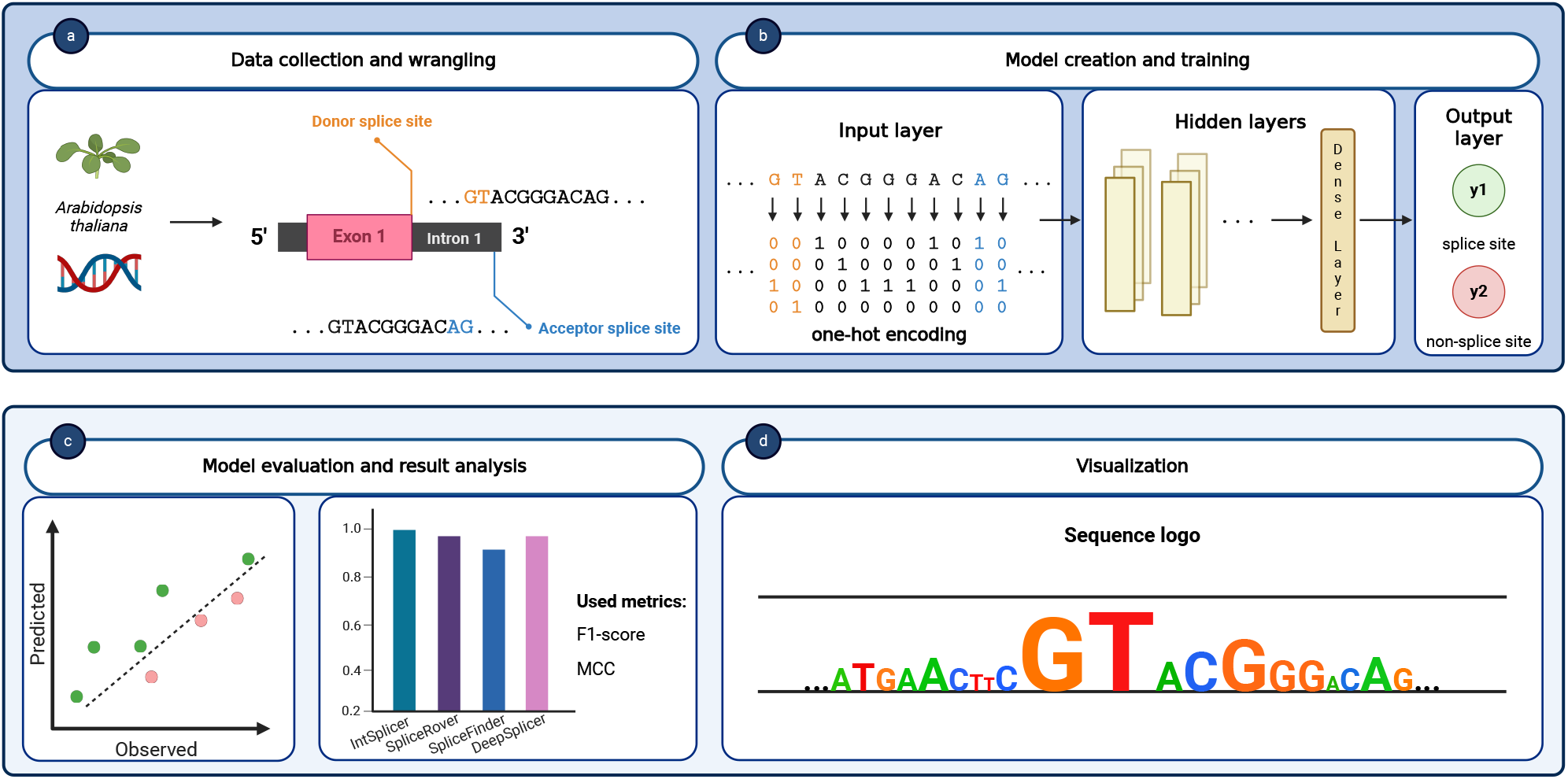
Study workflow. Our study followed a structured workflow, starting with the collection and identification of U2-type intron data from *Arabidopsis thaliana*, which was then categorized based on the hypotheses put forward (see panel **(a)**). The next step involved preprocessing the data, converting it into a one-hot encoded format suitable for input into the deep learning models (see panel **(b)**). We then proceeded to evaluate our models and analyzed the results obtained, using two metrics, F1-score and MCC, as shown in panel **(c)**. Finally, we focused on the visualization of our findings, using sequence logos, as shown in panel **(d)**.

#### 1. IntSplicer

Our model consists of multiple convolutional layers that play a crucial role in capturing essential sequence features, which are specific motifs that are crucial for identifying splice sites. The first layer consists of 64 filters with a kernel size of 10 and a stride of 4, followed by subsequent layers with increasing filter sizes (128, 256, and 512) and varying kernel sizes (3 and 2). Rectified Linear Unit (ReLU) activation functions are applied to introduce non-linearity. Between convolutional layers, max-pooling layers with a pool size of 2 are incorporated to down-sample the feature maps and reduce dimensionality. To prevent overfitting, dropout layers are placed after each max-pooling layer. Following the convolutional layers, a flatten layer transforms the feature maps into a one-dimensional vector. Next, a dense layer with 512 neurons and ReLU activation provides high-level abstractions of the extracted features. The final layer, consisting of two neurons with softmax activation, performs the binary classification for splice site prediction. IntSplicer, with its 1,962,946 trainable parameters, stands as a model of moderate complexity when compared to its counterparts.

#### 2. SpliceRover [11]

SpliceRover focuses on human and *Arabidopsis thaliana* data. This CNN model uses convolutional layers with ReLU activations, dropout layers, and a final 512-unit dense layer leading to a softmax binary classification layer. The architecture and approach of this model offer a substantial benchmark in terms of splice site classification effectiveness. SpliceRover has the highest number of trainable parameters (12,645,778) among the models compared. This suggests a considerably more complex model architecture, potentially allowing for more nuanced learning and pattern recognition capabilities. However, such complexity could also lead to challenges like overfitting, especially if the training data are not sufficiently large or diverse. Compared to SpliceRover, IntSplicer is substantially less complex, with roughly 85% fewer trainable parameters. This could imply a more streamlined or focused learning approach, resulting in faster training times and reduced computational resource requirements.

#### 3. SpliceFinder [12]

SpliceFinder consists of a single convolutional layer, followed by one fully-connected layer and a softmax layer for classification. Its structure, emphasizing the capture of local sequence patterns and complex pattern processing, served as a key comparative model for evaluating the efficiency and effectiveness of our CNN model in splice site detection. The parameter count of SpliceFinder (2,012,152) is slightly higher than the parameter count of IntSplicer but still operates on a similar scale. The closeness in the number of parameters suggests that both models may have similar complexities and potentially similar capabilities in terms of learning and generalization.

#### 4. DeepSplicer [14]

DeepSplicer, another CNN-based architecture, includes three convolutional layers, alongside flatten, fully-connected, dropout, and softmax layers. DeepSplicer has a comparable number of trainable parameters (2,057,252) to SpliceFinder and slightly more than IntSplicer. This similarity in model size suggests that DeepSplicer and IntSplicer might have similar computational demands and generalization capabilities.

### Experimental setup

For training the models, we adopted a standardized approach to hyperparameter selection to ensure comparability and consistency across all models. This approach involved utilizing the Adam optimizer with a learning rate of 0.001, the categorical cross-entropy loss function, and a batch size of 64, which empirically demonstrated optimal effectiveness in tests compared to batch sizes of 32, 128, and 256.

The Adam optimizer was selected for its effectiveness in handling sparse gradients and its adaptive learning rate properties, making it particularly well-suited for the models. The choice of a 0.001 learning rate was based on its widespread adoption as a value that balances fast convergence with the stability of the training process across a variety of tasks and datasets.

The uniform application of the different hyperparameters across all models helped to facilitate a fair and direct comparison of their effectiveness, removing potential biases that could arise from different training conditions.

We used a 5-fold cross-validation strategy with stratification to maintain the proportion of classes across folds, ensuring a robust evaluation framework. Considering that we organized the SpliceMachine dataset into six distinct subsets for each of the two hypotheses, making a total of 12 datasets, and that we applied 5-fold cross-validation to each, we conducted a total of 30 training sessions per hypothesis. The training sessions were performed on the training set portion of each dataset.

To further enhance the effectiveness of the training process and prevent overfitting, we used early-stopping. This technique monitors the validation loss, stopping the training if no improvement was observed after a certain number of epochs, thereby ensuring that the models were not over-trained. We set the total number of epochs to 30 and used early-stopping with patience 5, which means that training stops if no improvement is observed on the validation loss after 5 epochs.

## Results

In the context of splice site prediction, True Positives (TP) are splice sites correctly identified as either donor or acceptor sites. True Negatives (TN) are non-splice sites correctly identified as not being either donor or acceptor sites. False Positives (FP) are non-splice sites incorrectly identified as either donor or acceptor sites, and False Negatives (FN) are splice sites incorrectly identified as not being either donor or acceptor sites. With this understanding, we used two evaluation metrics to assess the prediction effectiveness of the models:

1. F1-score: The F1-score is a statistical measure used to evaluate the accuracy of a test. It considers both the precision (the proportion of true positive results in all positive predictions) and the recall (the proportion of true positive results over all actual positives). Specifically, the F1-score is the harmonic mean of precision and recall, calculated using the following formula:

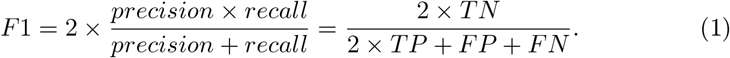
2. Matthews Correlation Coefficient (MCC): The MCC is a correlation coefficient used to measure the quality of binary classifications. It takes into account true and false positives and negatives, providing a score between *−*1 and +1. A score of +1 indicates perfect prediction, 0 no better than random prediction, and *−*1 total disagreement between prediction and observation. MCC is considered a balanced measure that can be used even if the classes are of very different sizes. MCC is calculated using the following formula:

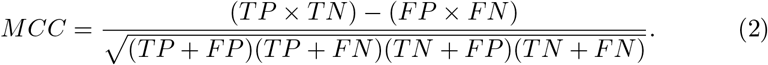

The evaluation and visualization were performed on the test set of each of the 12 subsets, ensuring that our assessment reflected the ability of the models to generalize on unseen data. Since during training we applied 5-fold cross-validation for each of the subsets based on the two hypotheses, we calculated the F1-score and MCC for each fold. Then, we used their average and standard deviation to evaluate the effectiveness.

### Short versus long U2-type introns

We refer to the datasets based on U2-type intron lengths as (1) the short dataset, consisting of DNA sequences with short U2-type introns, (2) the long dataset, consisting of DNA sequences with long U2-type introns, and (3) the length-mixed dataset, containing sequences with an equal mixture of short and long U2-type introns.

As shown in Table 3, for donor splice sites, the comparison of effectiveness across different datasets indicates that models generally achieve a slightly higher average F1-score and average MCC for the short dataset compared to the long and length-mixed datasets. While the effectiveness among the models is competitive, DeepSplicer shows a marginal advantage in both average F1-score and average MCC on the short dataset. On the other hand, the SpliceFinder model obtains the lowest average F1-score and average MCC on the long dataset.

**Table 3.** Average F1-score and average MCC for donor splice sites.

|  |  | IntSplicer | SpliceRover | SpliceFinder | DeepSplicer |
| --- | --- | --- | --- | --- | --- |
| Short | F1-score | *0.9456±0.0031* | *0.9435±0.0037* | *0.9263±0.0038* | **0.9463±0.0023** |
|  | MCC | *0.9347±0.0037* | *0.9322±0.0044* | *0.9119±0.0043* | **0.9357±0.0028** |
| Long | F1-score | 0.9398±0.0045 | 0.9411±0.0048 | 0.9097±0.0052 | 0.9401±0.0032 |
|  | MCC | 0.9278±0.0053 | 0.9294±0.0057 | 0.8918±0.0062 | 0.9282±0.0038 |
| Length-mixed | F1-score | 0.9318±0.0043 | 0.9298±0.0032 | 0.9120±0.0023 | 0.9387±0.0090 |
|  | MCC | 0.9182±0.0051 | 0.9157±0.0039 | 0.8947±0.0030 | 0.9266±0.0107 |

Summary of the effectiveness of the different donor splice site prediction models for the short, long, and length-mixed datasets. Results in italic indicate the highest effectiveness among the datasets, and results in bold indicate the highest effectiveness among the models.

As shown in Fig 5, for donor splice sites, IntSplicer performs better on the short dataset, achieving a higher median F1-score and median MCC, and exhibiting less variability compared to the long and the length-mixed datasets. SpliceRover has similar median F1-scores and median MCCs for the short and long datasets but a relatively low median F1-score and median MCC for the length-mixed dataset. SpliceFinder has the lowest median F1-score and median MCC among all the models. For DeepSplicer, the median F1-score and median MCC are comparable to those of the IntSplicer and SpliceRover models for the short dataset. However, it is also worth noting that the median F1-scores and median MCCs obtained by DeepSplicer and SpliceFinder for the length-mixed dataset are slightly higher compared to those obtained for the long dataset.

**Fig 5.**
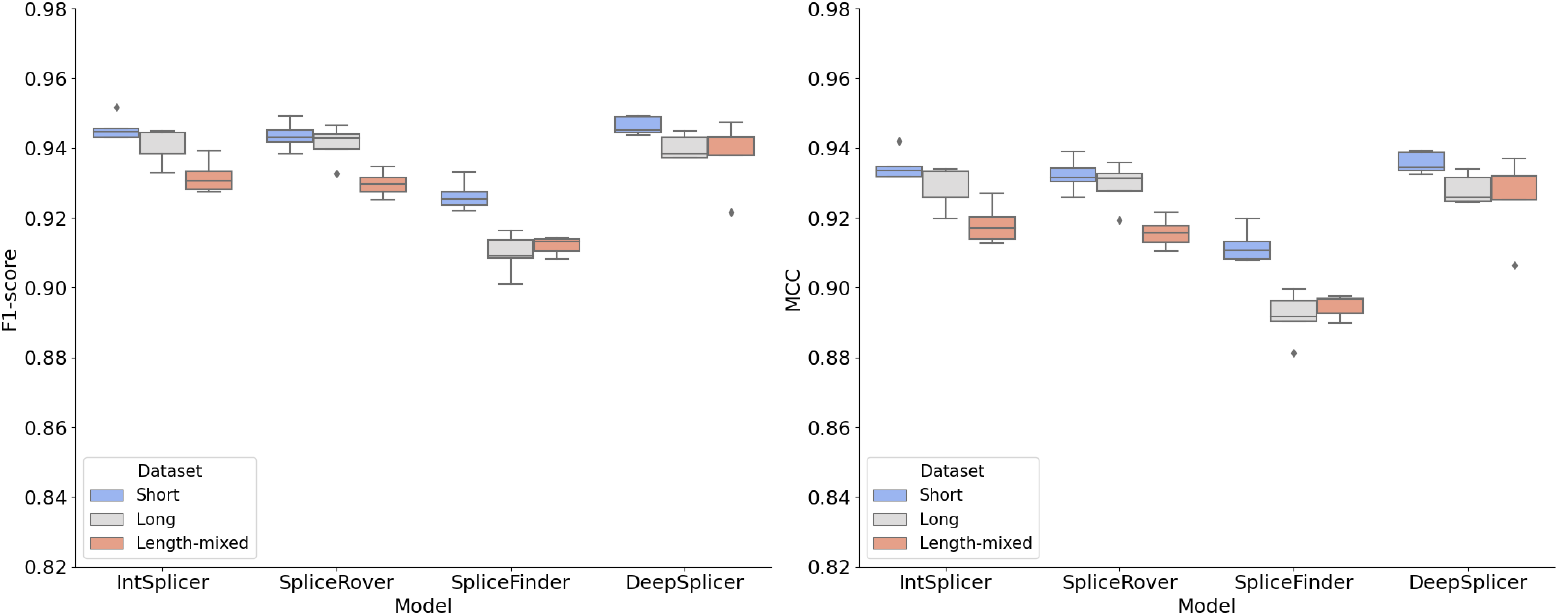
Comparison of model results for donor splice sites. F1-score distribution (left panel) and MCC distribution (right panel) from 5-fold cross-validation test results on the short, long, and length-mixed datasets.

Results for the acceptor splice site are shown in Table 4. The models generally achieve higher average F1-scores and average MCCs on the short dataset. SpliceRover demonstrates a different tendency, with its effectiveness peaking on the long dataset in terms of both average F1-score and average MCC. DeepSplicer presents a slight edge in both metrics for the short dataset compared to the other models, whereas SpliceFinder has the lowest average F1-score and average MCC for the length-mixed dataset.

**Table 4.** Average F1-score and average MCC for acceptor splice sites.

|  |  | IntSplicer | SpliceRover | SpliceFinder | DeepSplicer |
| --- | --- | --- | --- | --- | --- |
| Short | F1-score | *0.9233±0.0029* | 0.9184±0.0083 | *0.9116±0.0056* | **0.9326±0.0082** |
|  | MCC | *0.9080±0.0034* | 0.9024±0.0097 | *0.8942±0.0067* | **0.9192±0.0095** |
| Long | F1-score | 0.9207±0.0044 | *0.9191±0.0046* | 0.8853±0.0044 | 0.9178±0.0059 |
|  | MCC | 0.9047±0.0053 | *0.9030±0.0055* | 0.8622±0.0052 | 0.9013±0.0070 |
| Length-mixed | F1-score | 0.9084±0.0048 | 0.8989±0.0030 | 0.8664±0.0059 | 0.8985±0.0045 |
|  | MCC | 0.8899±0.0058 | 0.8786±0.0036 | 0.8414±0.0055 | 0.8786±0.0052 |

Summary of the effectiveness of the different acceptor splice site models for the short, long, and length-mixed datasets. Results in italic indicate the highest effectiveness among the datasets, and results in bold indicate the highest effectiveness among the models.

As shown in Fig 6, for acceptor splice sites, DeepSplicer achieves the highest median F1-score and median MCC on the short dataset, while SpliceFinder has the lowest median F1-score and median MCC on the length-mixed dataset across all models. The median F1-scores and median MCCs obtained by IntSplicer and SpliceRover for the short dataset are comparable. IntSplicer, SpliceRover, and DeepSplicer have similar median F1-scores and median MCCs for the long dataset. Overall, each model achieves the highest effectiveness for the short dataset and the lowest effectiveness for the length-mixed dataset.

**Fig 6.**
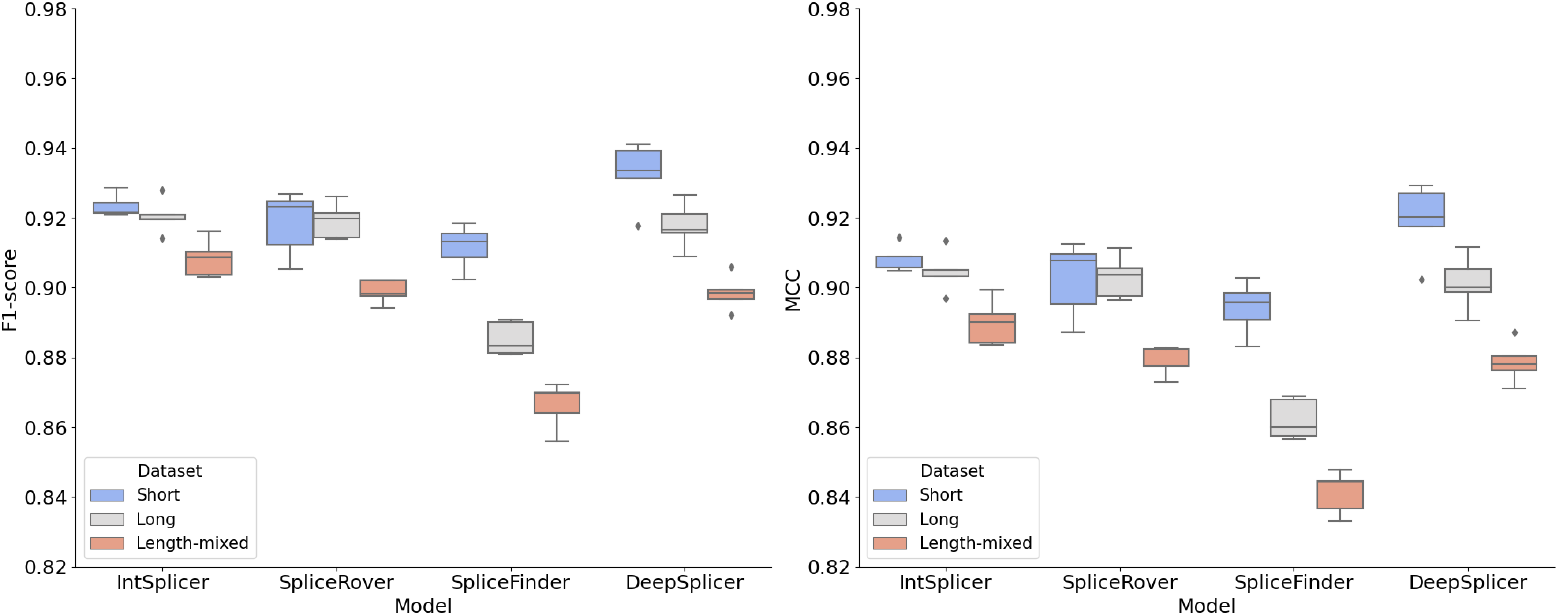
Comparison of model results for acceptor splice sites. F1-score distribution (left panel) and MCC distribution (right panel) from 5-fold cross-validation test results on the short, long, and length-mixed datasets.

### Single versus multiple U2-type introns

Considering the number of U2-type introns per sequence, we refer to the datasets used as (1) the single dataset, consisting of DNA sequences with one U2-type intron per sequence, (2) the multiple dataset, consisting of DNA sequences with more than one U2-type intron per sequence, and (3) the count-mixed dataset, containing sequences with an equal mixture of one and multiple U2-type introns per sequence.

Table 5 show the results for donor splice sites. We observe the highest average F1-score and average MCC on the multiple dataset across all the models, except for the SpliceFinder model. A comparatively lower effectiveness is obtained for the single and count-mixed datasets. Compared to the other models, SpliceRover achieves the highest average F1-score and average MCC on the multiple dataset, while SpliceFinder records the lowest average F1-score and average MCC on the single dataset.

**Table 5.** Average F1-score and MCC for donor splice sites.

|  |  | IntSplicer | SpliceRover | SpliceFinder | DeepSplicer |
| --- | --- | --- | --- | --- | --- |
| Single | F1-score | 0.9371±0.0049 | 0.9318±0.0011 | 0.9159±0.0049 | 0.9282±0.0023 |
| | MCC | 0.9246±0.0060 | 0.9182±0.0013 | 0.8990±0.0059 | 0.9140±0.0027 |
| Multiple | F1-score | *0.9503±0.0026* | **0.9543±0.0056** | 0.9224±0.0036 | *0.9426±0.0023* |
| | MCC | *0.9404±0.0032* | **0.9453±0.0067** | 0.9071±0.0044 | *0.9311±0.0027* |
| Count-mixed | F1-score | 0.9385±0.0025 | 0.9329±0.0065 | *0.9250±0.0005* | 0.9258±0.0080 |
| | MCC | 0.9264±0.0028 | 0.9196±0.0078 | *0.9104±0.0008* | 0.9110±0.0096 |

Summary of the effectiveness of the different donor splice site prediction models for the single, multiple, and count-mixed datasets. Results in italic indicate the highest effectiveness among the datasets, and results in bold indicate the highest effectiveness among the models.

As seen in Fig 7, IntSplicer and SpliceRover achieve comparable median F1-scores and median MCCs on the multiple dataset. Furthermore, SpliceFinder reaches its lowest median F1-score and median MCC for the single dataset. For IntSplicer, SpliceRover, and DeepSplicer, the median F1-scores and median MCCs are similar for the single and count-mixed datasets. Except for SpliceFinder, the models show a consistent trend, with all achieving higher median F1-scores and median MCCs for the multiple dataset.

**Fig 7.**
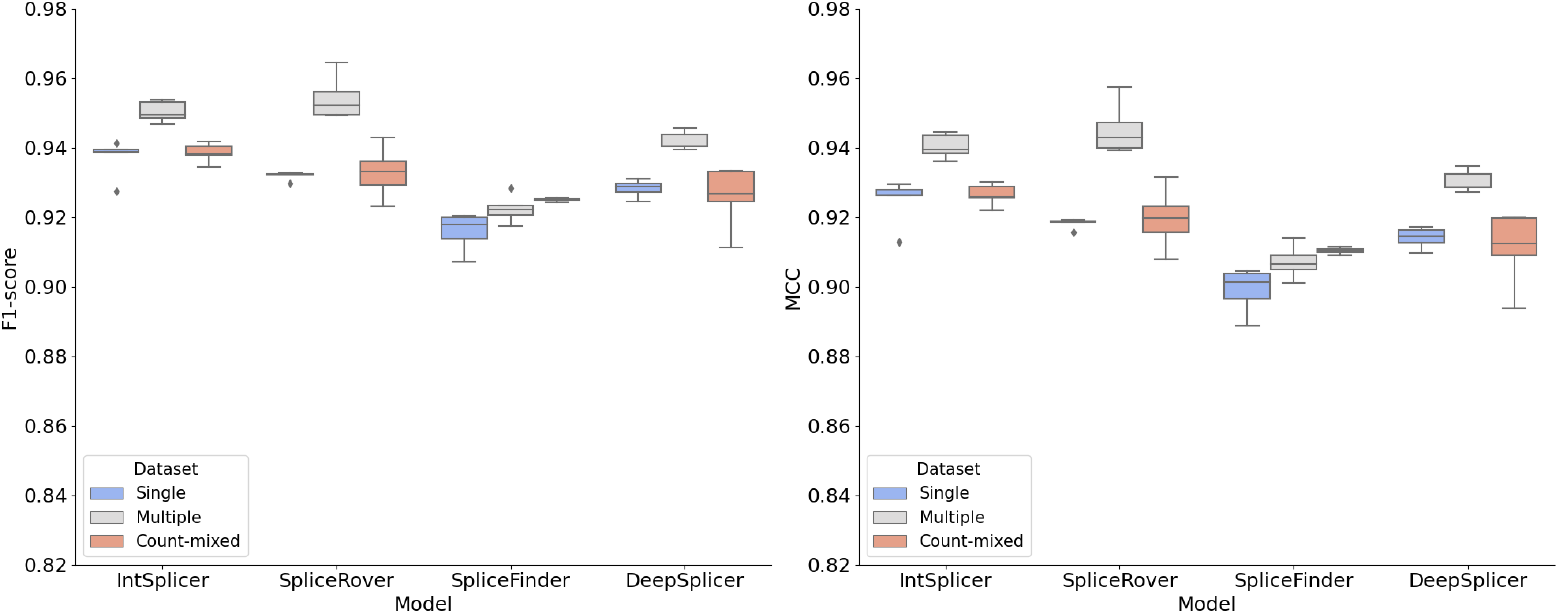
Comparison of model results for donor splice sites. F1-score distribution (left panel) and MCC distribution (right panel) from 5-fold cross-validation test results on the single, multiple, and count-mixed datasets.

In Table 6, we observe that for acceptor splice sites, each model achieves the highest average F1-score and average MCC on the multiple dataset, with IntSplicer having the highest effectiveness. SpliceFinder obtains the lowest average F1-score and average MCC among all the models on the single dataset. Unlike SpliceFinder which achieves an average F1-score and average MCC on the count-mixed dataset that is higher than those achieved on the single dataset, the other models obtain their lowest average F1-scores and average MCCs on the count-mixed dataset.

**Table 6.** Average F1-score and MCC for acceptor splice sites.

|  |  | IntSplicer | SpliceRover | SpliceFinder | DeepSplicer |
| --- | --- | --- | --- | --- | --- |
| Single | F1-score | $0.9158 \pm 0.0051$ | $0.9000 \pm 0.0060$ | $0.8803 \pm 0.0075$ | $0.9055 \pm 0.0055$ |
| | MCC | $0.8994 \pm 0.0063$ | $0.8806 \pm 0.0070$ | $0.8564 \pm 0.0089$ | $0.8872 \pm 0.0065$ |
| Multiple | F1-score | **$0.9187 \pm 0.0070$** | *$0.9176 \pm 0.0054$* | *$0.9004 \pm 0.0035$* | *$0.9098 \pm 0.0060$* |
|  | MCC | **$0.9028 \pm 0.0085$** | *$0.9013 \pm 0.0064$* | *$0.8811 \pm 0.0041$* | *$0.8931 \pm 0.0070$* |
| Count-mixed | F1-score | $0.9137 \pm 0.0024$ | $0.8866 \pm 0.0099$ | $0.8839 \pm 0.0052$ | $0.8985 \pm 0.0047$ |
| | MCC | $0.8964 \pm 0.0029$ | $0.8647 \pm 0.0110$ | $0.8610 \pm 0.0061$ | $0.8787 \pm 0.0057$ |

Summary of the effectiveness of the different acceptor splice site prediction models for the single, multiple, and count-mixed datasets. Results in italic indicate the highest effectiveness among the datasets, and results in bold indicate the highest effectiveness among the models.

As shown in Fig 8, IntSplicer and SpliceRover have the highest and comparable median F1-scores and median MCCs for the multiple dataset. While IntSplicer and SpliceFinder also show similar median F1-scores and median MCCs for the single and count-mixed datasets, we observe that the effectiveness of SpliceFinder is the lowest among all models. For DeepSplicer, the median F1-score and median MCC remain consistent when comparing the single and multiple datasets.

**Fig 8.**
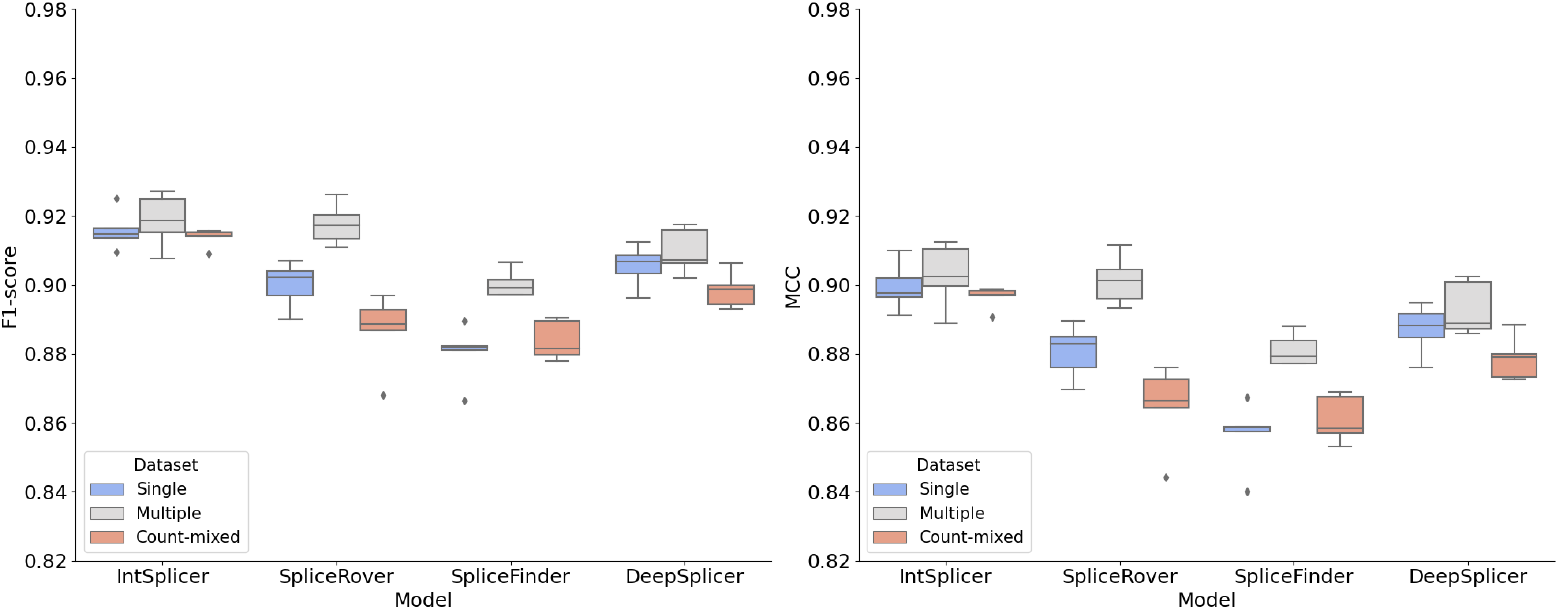
Comparison of model results for acceptor splice sites. F1-score distribution (left panel) and MCC (right panel) from 5-fold cross-validation test results on the single, multiple, and count-mixed datasets.

### Visualization

For visualisation purposes, we applied the saliency maps method described in [30] to calculate the contribution scores for each nucleotide in a DNA sequence with a true positive splice site. This method involves computing the gradients of the output of the model with respect to its input, known as saliency scores. After calculating the saliency scores, we normalized them for better visualization, as implemented in [11]. This normalization process can be mathematically represented as follows:

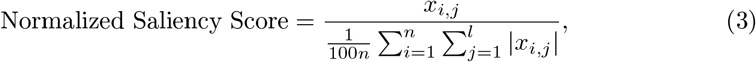

where *x*_*i,j*_ represents the contribution score for the *j*-th position of the *i*-th sequence, *n* is the total number of sequences, *l* is the sequence length, and the denominator computes the average of the absolute score sums across all sequences, scaled by 100. For example, for the sequence ACTG with scores (0.1, *−*0.2, 0.3, *−*0.05) and the sequence CGTG with scores (0.5, 0.6, 0.15, *−*0.01), the absolute sums are computed as 0.65 and 1.26, respectively. The average of these sums is then calculated, resulting in 0.955. This average is scaled down by a factor of 100, yielding a normalization constant *S*_avg_ = 0.00955. Using this constant, each score is normalized by dividing it by *S*_avg_. The resulting normalized scores for the sequence `ACTG` are approximately (10.47, *−*20.94, 31.41, *−*5.24) and for `CGTG` are (52.36, 62.83, 15.71, *−*1.05). This approach ensures that the scores are adjusted relative to the average magnitude of scores across all sequences, enabling a consistent scale for comparison. The normalized scores are then associated with each nucleotide in the DNA sequence, creating a saliency map [31]. We used these saliency scores, also called contribution scores, to generate high-quality sequence logos using the Python package Logomaker [32].

Fig 9 show the sequence logos for donor splice sites for the short dataset (top sequence logo), the long dataset (middle sequence logo), and the length-mixed dataset (bottom sequence logo). These logos were generated for DeepSplicer, which achieved the highest average F1-score and average MCC for donor splice sites according to the first hypothesis. The donor splice sites are positioned at 0 and 1. The U2-intron length is approximately 80 bp in the short dataset and about 105 bp in the long dataset. The length-mixed dataset displays a distinctive motif, including occurrences of the `G` nucleotide with a positive contribution score near position 30, around 60, and between 60 and 70. Additionally, for the length-mixed dataset, there are multiple `AG` dinucleotides between positions 80 and 90, all of which are candidate acceptor splice sites, indicating variable lengths of the U2-type introns. Similarly, Fig 10 presents the sequence logos for acceptor splice sites for the short dataset (top sequence logo), the long dataset (middle sequence logo), and the length-mixed dataset (bottom sequence logo). DeepSplicer was also used for these logos due to its highest average F1-score and average MCC among models tested for acceptor splice sites under the first hypothesis. The acceptor splice sites are located at positions 0 and 1. In the short dataset, the U2-type intron covers about 76 base pairs from position *−*75 to position 1. In the long dataset, the U2-type intron is of approximately 96 base pairs from position *−*95 to position 1. In the length-mixed dataset, a `GT` dinucleotide appears at positions *−*85 and *−*77, both potential candidate donor splice sites.

**Fig 9.**
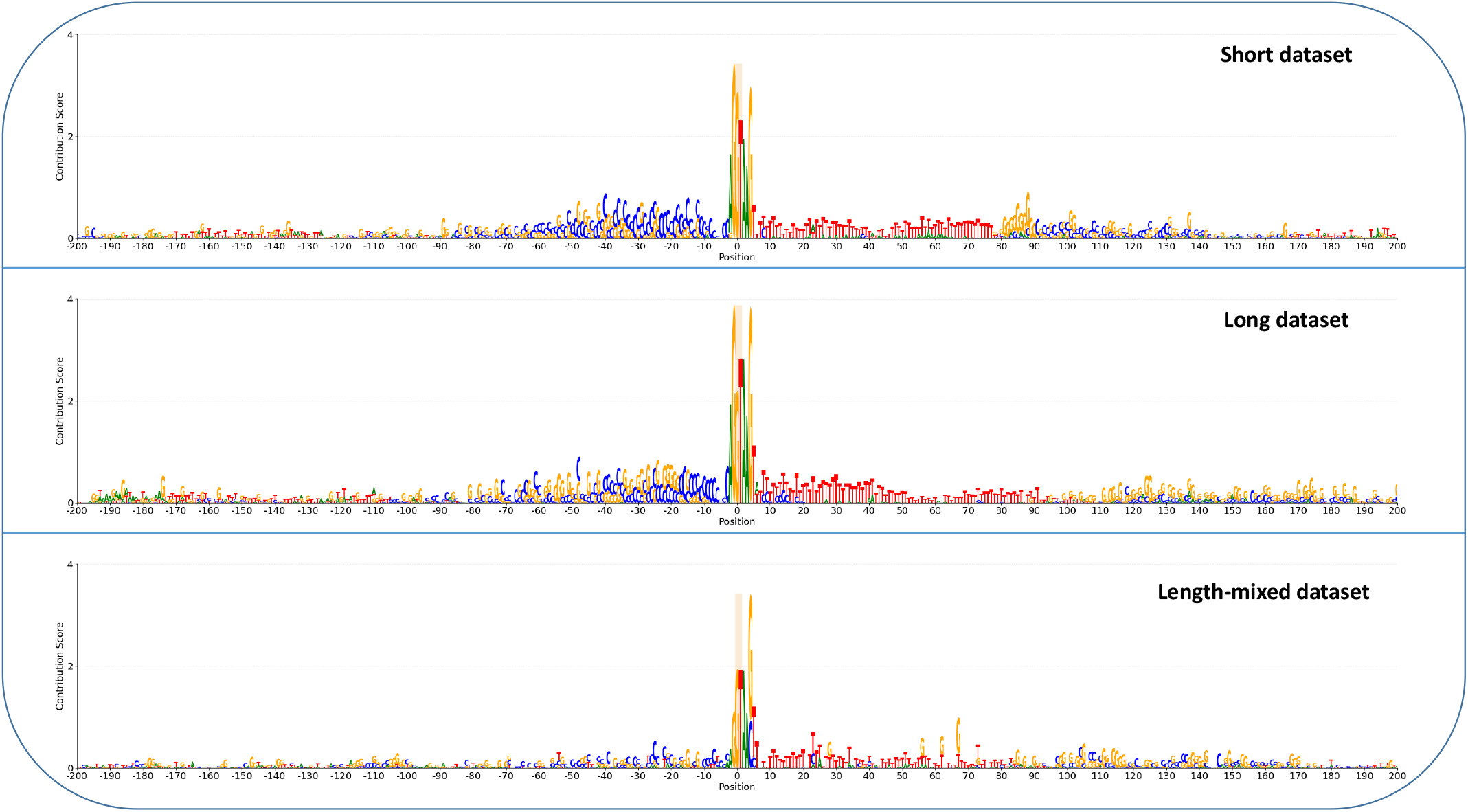
Sequence logos for donor splice sites based on U2-type intron length. Comparison of sequence logos for the short dataset (top), for the long dataset (middle), and for the length-mixed dataset (bottom).

**Fig 10.**
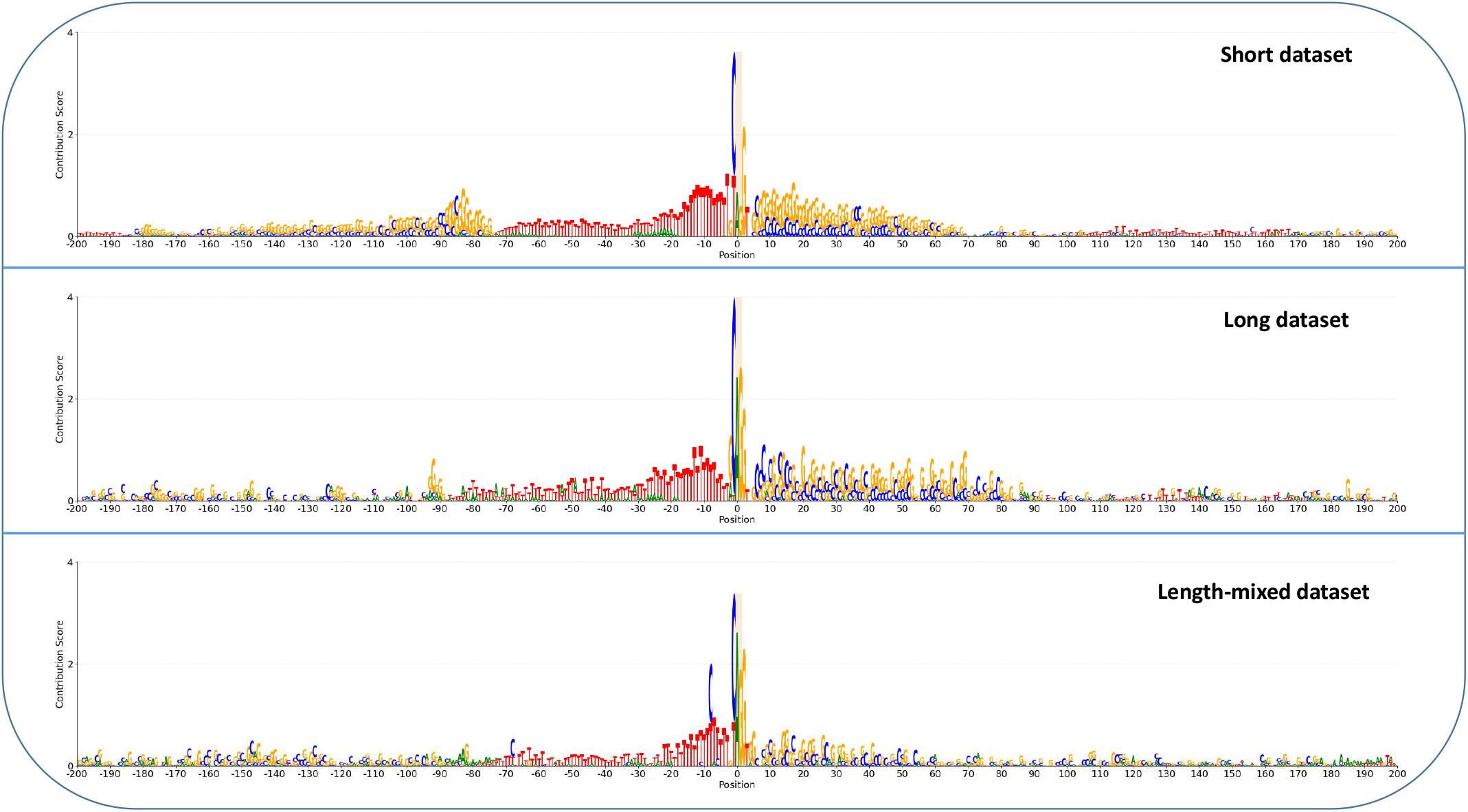
Sequence logos for acceptor splice sites based on U2-type intron length. Comparison of sequence logos for the short dataset (top), for the long dataset (middle), and for the length-mixed dataset (bottom).

Fig 11 shows the sequence logos for donor splice sites for the single dataset (top sequence logo), the multiple dataset (middle sequence logo), and the count-mixed dataset (bottom sequence logo), generated for SpliceRover, as it achieved the highest average F1-score and average MCC for donor splice sites for the second hypothesis. Donor splice sites are located at positions 0 and 1. In the single dataset, the U2-intron is seen from position 0 to around position 100. The multiple dataset shows a U2-type intron between positions *−*155 and *−*75, and another from position 0 to 91, illustrating that the models can identify two different U2-type introns within sequences. For the count-mixed dataset, the U2-type intron from position 0 to around position 95 is visible in the sequence logo. Similarly, Fig 12 shows the sequence logos for acceptor splice sites for the single dataset (top sequence logo), multiple dataset (middle sequence logo), and count-mixed dataset (bottom sequence logo), generated using IntSplicer, which also achieved the highest average F1-score and average MCC among the models for acceptor splice sites under the second hypothesis. The acceptor splice sites are located at positions 0 and 1. For the single dataset, the U2-type intron is located from position *−*90 to position 1. In the multiple dataset, there are two distinct U2-type introns, one located from position *−*85 to 1, and another from position 70 to 169. In the count-mixed dataset, there is a U2-type intron from position *−*82 to 1, and another region from position 100 to 170 with a very low contribution score. These sequence logos demonstrate the models’ sensitivity to the number of U2-type introns within sequences. These visualizations effectively illustrate how U2-type introns’ length and count contribute to the prediction effectiveness of the models.

**Fig 11.**
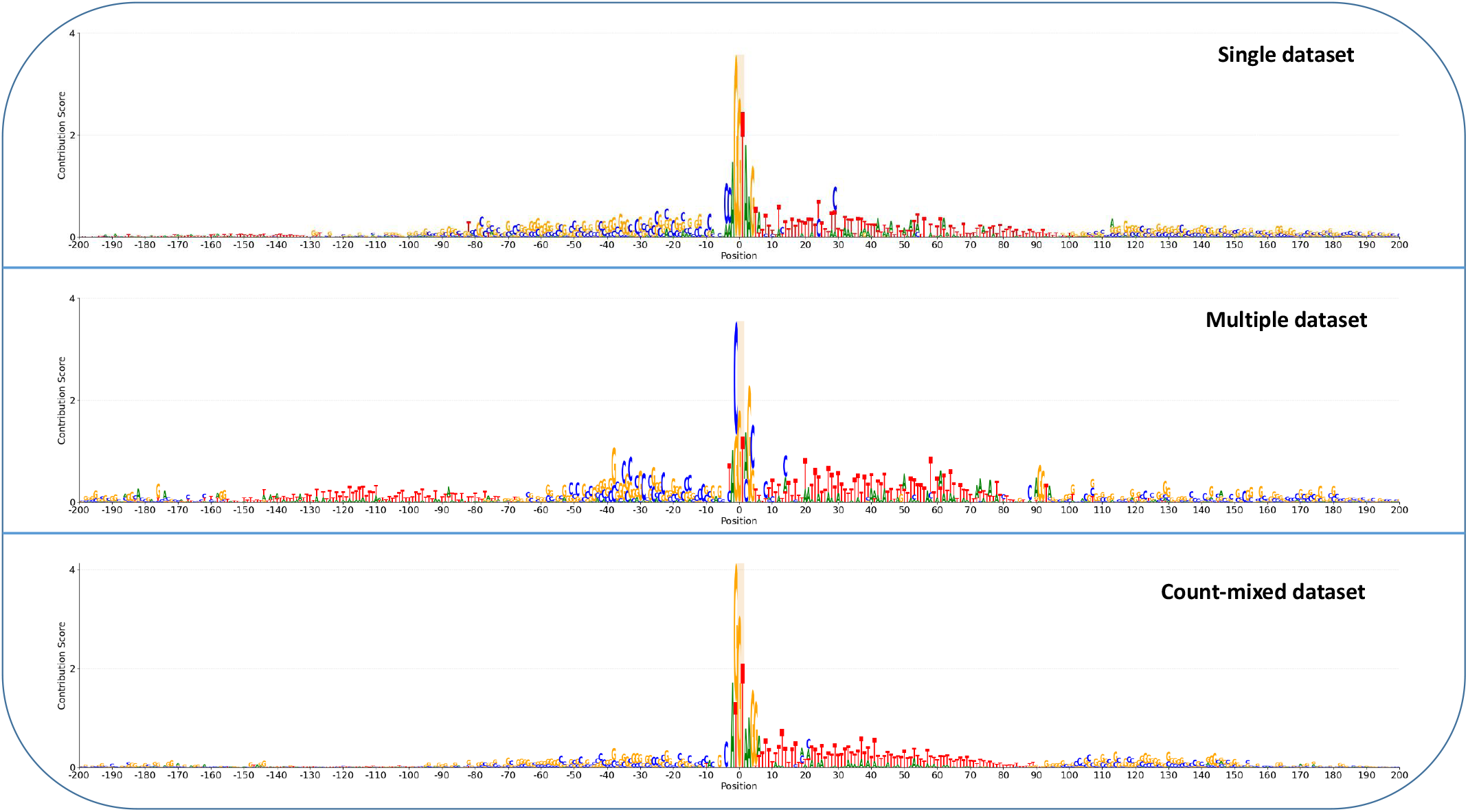
Sequence logos for donor splice sites based on U2-type intron count. Comparison of sequence logos for the single dataset (top), for the multiple dataset (middle), and for the count-mixed dataset (bottom).

**Fig 12.**
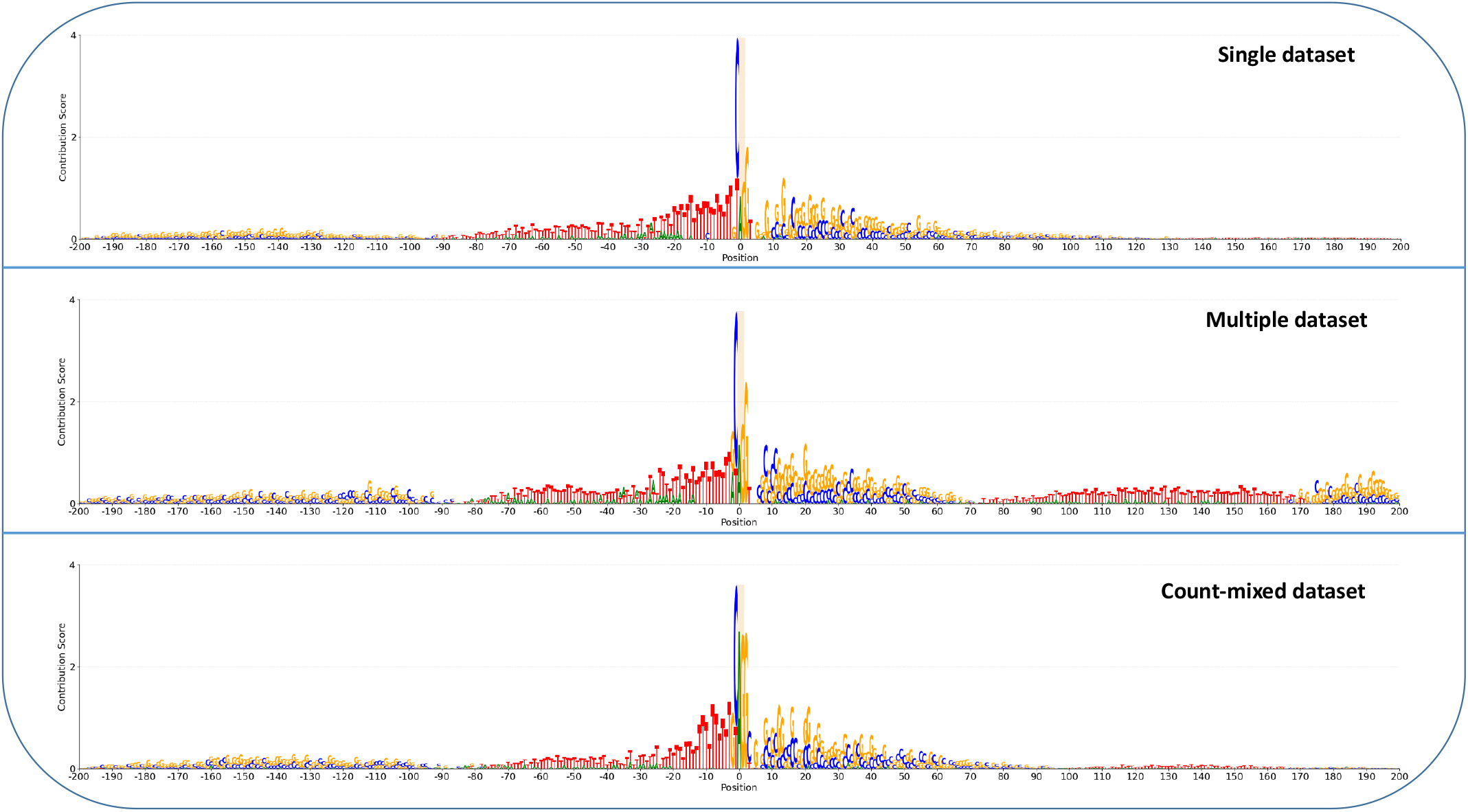
Sequence logos for acceptor splice sites based on U2-type intron count. Comparison of sequence logos for the single dataset (top), for the multiple dataset (middle), and for the count-mixed dataset (bottom).

## Discussion

### Models trained on datasets with short U2-type introns demonstrate improved predictive effectiveness

The first hypothesis suggested that models trained on datasets with short U2-type introns would outperform those trained on datasets with long U2-type introns in predicting both donor and acceptor splice sites. Our experimental results confirmed this hypothesis for donor splice site prediction across all the models and for acceptor splice site prediction for all the models except SpliceRover. The underlying premise is that longer introns may introduce spatial complexities and a higher likelihood of engaging non-canonical splice sites [23]. Such complexities could undermine the precision of the splice site predictions made by the different models, hence the emphasis on shorter introns which tend to be simpler and are less susceptible to misinterpreting exon-skipping events [24]. Furthermore, long introns could also contain more control signals like enhancers or silencers, affecting which site gets chosen for splicing. These elements in the introns are important for adjusting where splicing happens [33].

Correspondingly, our experimental results align with the observations in [34], which proposed a linkage between short intron lengths and a more uniform splicing mechanism. This uniformity might arise from an increased density of splicing signals in shorter introns. Additionally, according to [35, 36], shorter introns may lead to more effective splicing machinery processing and recognition, which in turn could facilitate more consistent regulation of gene expression.

The biological implications of these findings reinforce the concept that intron length may be a critical element in the precision of splicing. The observed patterns suggest that shorter introns enhance the reliability of gene expression regulation, highlighting their potential evolutionary advantage in maintaining genomic integrity.

### Having multiple U2-type introns per sequence improves splice site prediction effectiveness

The second hypothesis dealt with the impact of intron multiplicity within sequences. Our experimental results pointed to higher effectiveness in splice site prediction for datasets containing multiple U2-type introns per sequence. This observation is consistent with the underlying biological mechanisms governing RNA splicing, where the presence of multiple introns within a gene sequence has been shown to positively impact splicing kinetics and gene expression regulation [28].

Kinetic analyses of splicing have illuminated that the excision of the first intron can substantially improve the effectiveness of subsequent intron removals, potentially due to a priming effect that facilitates the assembly of the splicing machinery or the recruitment of the EJC to the spliced mRNA. This mechanism suggests a synergistic effect, where each splicing event incrementally improves the precision of subsequent splicing within the same transcript.

Experimental evidence, such as studies on the ERECTA gene in *Arabidopsis thaliana*, further underscores the beneficial impact of multiple introns on gene expression. These studies demonstrate that while no single intron may be crucial for expression, the cumulative presence of multiple introns can substantially enhance mRNA accumulation and gene expression in an additive fashion [37]. This effect can also extend to the regulation of splicing factor abundance, indirectly influencing splice site selection accuracy.

These findings imply that multiple introns within a sequence may have a cumulative effect on enhancing the regulatory complexity and efficiency of gene expression [38], highlighting the biological importance of intron multiplicity in the context of genetic regulation.

## Conclusions

This study explored the complexity of splice site prediction in *Arabidopsis thaliana*, focusing on the impact of U2-type intron characteristics on the effectiveness of deep learning models. Through the investigation of two hypotheses, we explored how intron length and the number of introns per sequence influence the effectiveness of splice site prediction. Regarding the intron length, our findings show that the models trained and tested on datasets with shorter U2-type introns achieve higher prediction effectiveness compared to those trained and tested on dataset with long U2-type introns, for both donor and acceptor splice prediction. Regarding the number of introns per sequence, our results show that deep learning models trained and tested on datasets having multiple U2-type introns per sequence generally achieve higher prediction effectiveness compared to when trained and tested on datasets having a single U2-type intron per sequence. These results underscore the complexity of the splicing process, highlighting the importance of intron characteristics in the regulation of gene expression and demonstrating the need for advanced approaches capable of capturing the nuanced biological signals governing splice site selection. Future research will aim at integrating additional genomic features in our current methodology, broadening the scope of the data to include various intron types and configurations across different organisms. This could reveal insights into the universality or specificity of splicing mechanisms.

## Acknowledgments

The research activities described in this paper were funded by Ghent University Global Campus and Ghent University.

## Data availability

The dataset used in this work is available at https://zenodo.org/records/11178747. All annotated U2-type introns used in this study can be downloaded from https://introndb.lerner.ccf.org/static/fasta/TAIR10_U2.fasta. To ensure reproducibility, the code for this work can be found at https://github.com/EspoirKabanga/Impact_of_U2type_Introns_on_Splice_Site_Prediction.

## Notes

### Competing Interest Statement

The authors have declared no competing interest.

